# A Likelihood Ratio Test for Hybridization Under the Multispecies Coalescent

**DOI:** 10.1101/2023.06.20.545699

**Authors:** Jing Peng, Sungsik Kong, Laura Kubatko

## Abstract

Several methods have been developed to carry out a statistical test for hybridization at the species level, including the ABBA-BABA test and *HyDe*. Here, we propose a new method for detecting hybridization and quantifying the extent of hybridization. Our test computes the likelihood of a species tree that is possibly subject to hybridization using site pattern frequencies from genomic-scale datasets under the multispecies coalescent. To do this, we extend the calculation of the likelihood for site pattern frequency data for the 4-taxon symmetric and asymmetric species trees proposed in Chifman and Kubatko (2015) by incorporating an inheritance parameter, resulting in efficient computation of the likelihood under a scenario of hybridization. We use this likelihood computation to construct a likelihood ratio test that a given species is a hybrid of two parental species. Simulations demonstrate that our test is more powerful than existing tests of hybridization, including *HyDe*, and that it achieves the desired type I error rate. We apply the method to two empirical data sets, one for which hybridization is believed to have occurred and one for which previous methods have failed to detect hybridization.

## 1 Introduction

The deluge of genomic data available for phylogenetic study has confirmed the ubiquity of interspecific hybridization across the tree of life. Processes such as hybridization are ideally represented by phylogenetic networks, which generalize phylogenetic trees to include reticulate evolutionary events that allow the possibility that the ancestry of a species is derived from two (or more) independent evolutionary lineages. Phylogenetic networks can thus be used to model processes such as hybridization, horizontal gene transfer, gene duplication and loss, and recombination (Linder, 2004; Nakhleh, 2010). Despite the development of innovative inference algorithms (e.g., Solís-Lemus and Ané, 2016; Than *et al*., 2008), estimating a phylogenetic network is a challenging task because the methods currently available often scale poorly and are presently limited to the analysis of relatively small data sets (Hejase and Liu, 2016). An alternative approach is to employ methods that detect hybrids among a large number of genomic sequences, rather than attempting to estimate the phylogenetic network directly.

Model-based population genetic clustering approaches are widely used to identify hybrid individuals from genetic data, and serve as an important tool for understanding patterns of extant genetic variation. Often implemented within the maximum likelihood (Alexander *et al*., 2009) or Bayesian frameworks (Pritchard *et al*., 2000), these methods estimate contributions from a user-designated number of ancestral genetic pools (typically denoted by *κ*) to an individual’s ancestry through estimation of probabilistic quantities called ancestry coefficients. Despite the popularity of these methods in studies of hybridization on phylogenetic time scales, population clustering methods were not originally designed for this task but rather for the task of identifying population structure in contemporary populations. As a result, interpretation of the ancestry coefficients in a phylogenetic context is inevitably subjective and prone to mis- or over-interpretation of the historical processes, because different evolutionary scenarios can result in indistinguishable patterns (Barilani *et al*., 2007; Anderson and Dunham, 2008; Lawson *et al*., 2018). For example, gene flow is often conjectured to be responsible for an intermediate ancestry coefficient, even though incomplete lineage sorting (ILS) can lead to very similar patterns. These methods are also sensitive to the choice of markers, the level of genetic differentiation between populations, and the amount of data utilized (Vähä and Primmer, 2005; Latch *et al*., 2006; Kalinowski, 2011). The algorithms implemented in some of these tools involve unsupervised clustering, and stochastic simulations (e.g., *structure*, Pritchard *et al*., 2000) have demonstrated that these methods can produce different outcomes in replicate analyses due to label-switching or multimodality (Kopelman *et al*., 2015). While the former issue can be detected and eliminated through post-processing, the latter issue is much more difficult to assess. Therefore, we have recently recommended that the use of such methods be curtailed in favor of newer methods that are specifically designed to detect hybridization across phylogenetic time scales (Kong and Kubatko, 2021).

Simple and intuitive approaches based on site pattern frequencies have quickly gained popularity in detecting hybridization from genomic datasets. Some of the widely used methods include the *f*_3_ and *f*_4_ statistics (Reich *et al*., 2009) and Patterson’s *D*-statistic (Patterson *et al*., 2012), also known as the ABBA-BABA test (Green *et al*., 2010; Durand *et al*., 2011). Patterson’s *D*-statistic is used as the basis of a statistical test for which rejection of the null hypothesis indicates a history of introgression among the input taxa. Recent work in this area (e.g., Hibbins and Hahn (2019)) has included further development of the methodology in an attempt to quantify the direction and proportional contributions of the taxa involved in the introgression event. Kubatko and Chifman (2019) propose a coalescent-based method that uses phylogenetic invariants for detecting species that have arisen via hybridization, implemented in the computer program *HyDe* (Blischak *et al*., 2018). Unlike methods based on *D*-statistics, this method is not limited to the examination of a single individual per population and it has been shown to detect populations that may have arisen via hybrid speciation as well as their putative parental populations with statistical power that is similar to the *D*-statistic. In addition, *HyDe* estimates the inheritance parameter (*γ*) that quantifies the proportion of genomic contribution of each parental taxon to the hybrid species. Kong and Kubatko (2021) found that the accuracy of the *γ* estimates in *HyDe* is superior to the estimates of ancestry coefficient in population clustering methods, even when the amount of ILS is high, though large sample sizes may be required when extensive ILS is present. Another method that has been widely used to quantify *γ* is the *f*4-ratio statistic, although it has been found to be sensitive to violations of the underlying population model (Patterson *et al*., 2012).

In this paper we develop a method for detecting hybridization and quantifying the extent of hybridization by computing the likelihood of a species tree that is possibly subject to hybridization using site pattern frequencies from genomic-scale datasets under the multispecies coalescent. To do this, we extend the calculation of the likelihood for site pattern frequency data for the 4-taxon symmetric and asymmetric species trees proposed in Chifman and Kubatko (2015) by incorporating *γ*. Because our method uses site pattern frequencies calculated from either multi-locus data or single nucleotide polymorphism (SNP) data, the likelihood can be evaluated in a computationally efficient manner. We use these likelihood computations to construct a likelihood ratio test that a given species is a hybrid of two parental species. We use simulation to demonstrate that our test is more powerful than existing tests of hybridization, including *HyDe*, and that it achieves the desired type I error. We apply the method to two empirical data sets, one for which hybridization is believed to have occurred and one for which previous methods have failed to detect hybridization.

## 2 Methods

### 2.1 Likelihood of a 4-taxon network

Consider the rooted, 4-taxon network S in Figure 1, where the outgroup population is *O*, the two parental populations are *P* 1 and *P* 2, and *H* is the hybrid population with inheritance parameter *γ*. The species network S can be decomposed into S1 and S2, where the former can be obtained by removing the reticulation edge between *P* 1 and *H* and the latter can be obtained by removing the edge between *H* and *P* 2. In this case, sequences are assumed to evolve through gene trees, which arise either from species tree S1 with *H* as a sister taxon of *P* 2 with probability *γ*, or from S2 with *H* as a sister taxon of *P* 1 with probability 1 − *γ*. Note that we can summarize sequence data from the network S as site patterns.

**Figure 1:**
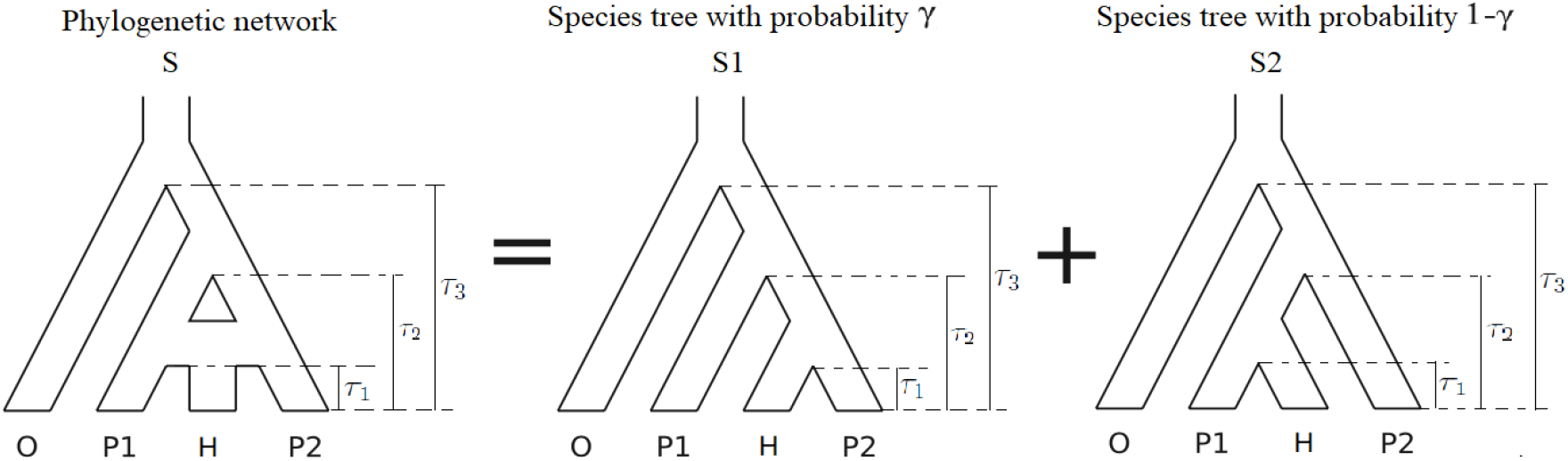
A rooted, 4-taxon network S with 3 branch length parameters *τ*_1_, *τ*_2_ and *τ*_3_. The inheritance parameter *γ* (and 1 *− γ*) can be used with the site pattern probabilities under species trees S1 and S2 to derive site pattern probabilities arising from the network.

For example, a site pattern AGCC represents a position in the alignment for which species *O, P* 1, *H* and *P* 2 have nucleotides A, G, C and C, respectively. In a 4-taxon network, there are 4^4^ = 256 possible site patterns.

To define the site pattern probabilities, consider a 4-taxon species tree with species *a, b, c*, and *d*, and let *i*_*j*_ ∈ {*A, C, G, T* } refer to the nucleotide observed for taxon *j* at the particular site under consideration. We refer to *i*_*a*_*i*_*b*_*i*_*c*_*i*_*d*_ as a site pattern for the species tree, and denote the probability of this site pattern by 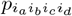. Chifman and Kubatko (2015) derived explicit expressions for the site pattern probabilities under the multispecies coalescent model with the JC69 (Jukes and Cantor, 1969) substitution model and the assumption of a constant effective population size parameter *θ* with branch lengths ***τ*** = (*τ*_1_, *τ*_2_, *τ*_3_) in coalescent units. Under this model and the molecular clock assumption, the rooted symmetric 4-leaf species tree ((*a, b*), (*c, d*)) will have 9 distinct site patterns probabilities:

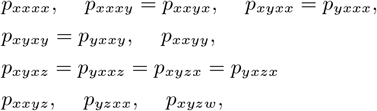

while the rooted asymmetric 4-leaf species tree (*a*, (*b*, (*c, d*))) will have 11 distinct site patterns probabilities:

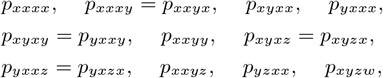

where *x, y, z* and *w* denote different nucleotide states. Therefore, for the network S in Figure 1, we will have 15 site pattern probabilities, with each one being a weighted average of the probabilities from species trees S1 and S2. For example, the probability of site pattern *xxxy* from the network S is:

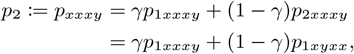

where *p*_1*xxxy*_ and *p*_2*xxxy*_ are the probabilities of site pattern *xxxy* from species trees S1 and S2, respectively. Similarly, we can write the other 14 site pattern probabilities:

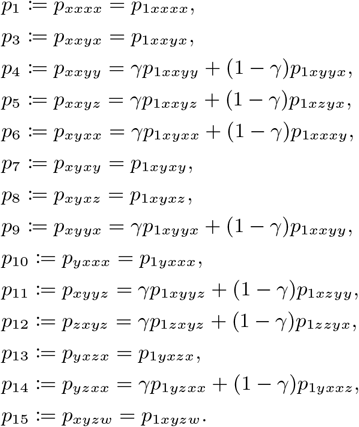

From the data, we observe the frequency with which site pattern *k, k* = 1, 2, …, 15 is observed, which we denote *d*_*k*_, in a sample of *n* sites. The entire data are then denoted by the vector *D* = (*d*_1_, …, *d*_15_).

The likelihood of network-associated parameters (*τ*_1_, *τ*_2_, *τ*_3_, *γ* and *θ*) given network topology S is then given by:

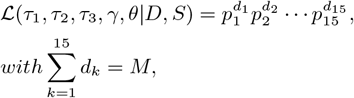

where *M* is the total number of sites.

### 2.2 Likelihood ratio test (LRT) of hybridization

Consider the 4-taxon species tree S2 in Figure 1, and note that a value of *γ* = 0 implies the absence of genetic contribution from *P* 2 to *H* following the divergence of *P* 2 from the ancestral species of *P* 1 and *H* (i.e., no hybridization). We can thus develop a formal statistical hypothesis test for hybridization between species *H* and *P* 2 by considering the following hypotheses:

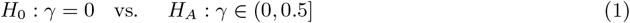

Rejection of the null hypothesis above indicates support for the reticulation event between *H* and *P* 2 for the tree S2, and *H* can be considered to be a hybrid of parental species *P* 1 and *P* 2. A failure to reject *H*_0_ implies that there is not sufficient evidence from the data to prefer the network S over the species tree S2.

We propose to test the hypotheses above using a likelihood ratio test, which has well-established statistical properties in general contexts (Wilks, 1938). The test statistic is given by

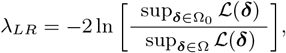

where ***δ*** is the vector of parameters, i.e.,

***δ*** = (*τ*_1_, *τ*_2_, *τ*_3_, *θ, γ*), and Ω is the parameter space for the network model. Specifically, Ω is a 5-dimensional space defined by *τ*_1_ ∈ (0, +∞), *τ*_2_ ∈ (*τ*_1_, *τ*_3_), *τ*_3_ ∈ (*τ*_2_, +∞), *θ* ∈ (0, +∞), *γ* ∈ [0, 0.5]. The space Ω_0_ is the subset of Ω defined by the null hypothesis, i.e., Ω_0_ is the 4-dimensional subspace obtained by fixing *γ* = 0. Standard statistical theory can be applied to see that the asymptotic distribution of *λ*_*LR*_ under the null hypothesis is a 50:50 mixture of a 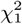 distribution and a point mass at 0 since the value of *γ* under the null hypothesis lies on the boundary of the parameter space (Self and Liang, 1987). Thus, the hypothesis test at level *α* can be carried out by comparing *λ*_*LR*_ with the 1 − *α* quantile of this mixture distribution.

We note here that to compute *λ*_*LR*_, we have to find the value of ***δ*** that maximizes the likelihood over both Ω and Ω_0_. Because this optimization procedure is constrained by the bounds on the branch lengths, the population size parameter and the hybridization parameter, we use the following reparameterization, to develop an efficient optimization algorithm:

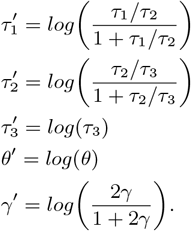

Defining ***δ***′, as the vector of transformed parameters, we must find the values of 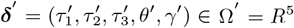 and 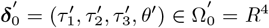 that maximize the likelihood. With this transformation, we perform a multidimensional optimization simultaneously for all parameters by applying the quasi-Newton method (BFGS) (Byrd *et al*., 1995; Fletcher and Reeves, 1964) for unconstrainned multi-dimensional optimization, which uses function values and gradients to search parameter space. The BFGS method has better computational complexity than Newton’s method, and because it uses an approximation of the gradient to carry out the search, it is expected to be more computationally efficient than gradient-free methods. However, because of correlation between *θ* and the *τ* s, the density is relatively flat for values of *θ*, which sometimes causes the BFGS optimization process to terminate prematurely. We have noticed that a crucial step in developing a good implementation of this method is the selection of a good starting point, especially for the population size parameter *θ*.

We thus obtain a starting point, *θ*_0_, by first setting a small lower bound (10^−5^ in our case), and then increasing it until at least one of the branch length moment estimates is larger than 0. Using this value as the upper bound, we then find an initial interval [*a, b*] for *θ*. This is a very wide interval, so we then implement a golden section search (Gill *et al*., 1981) to get a tighter interval for *θ*_0_. The disadvantage of golden section search is its slow convergence, so we set a large stopping tolerance, terminating the search once *b* − *a <* 0.01. Constrained in this updated interval [*a*_1_, *b*_1_], we finally use one-dimensional Brent optimization (Brent, 1973), and the optimal value is used as *θ*_0_. Brent optimization can achieve superlinear convergence via a combination of golden section and parabolic interpolation steps. This procedure was motivated by the work of Peng et al. (2022) (see their Supplemental Information for details).

For the branch lengths and the inheritance probability, we can simply use the moment estimators by adapting results in Kubatko et al. (2023) for networks. By solving the equations

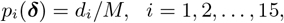

where *M* is the total number of sites, we obtain the following moment estimator for the branch lengths given *θ*_0_:

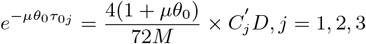

where

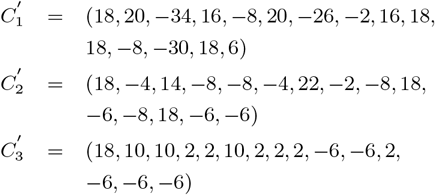

and *μ* is set to be 4/3 for the JC69 model. The moment estimator of *γ* can be obtained from any phylogenetic invariant; we use

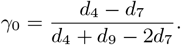

### 2.3 Simulation study

We first use simulations to assess the type I error and statistical power of testing hybridization using our method (LRT), *HyDe* and the ABBA-BABA test. Coalescent Independent Site (CIS) data is a natural fit for our method, because nucleotides are unlinked and assumed to arise from the coalescent model independently. Even though we are not generating CIS data in practice, these data are useful for verifying our method and theory under the correct model. To further simulate the performance of our method in application, we also implement this approach on multilocus data, in which all sites within a given locus are assumed to have evolved on the same genealogy and are not independent. A theoretical justification for this application can be found in Wascher and Kubatko (2021), which argues that methods developed for CIS data can also be applied to multilocus data, and we therefore consider both data types here.

To examine the performance of the three tests, we simulated two types of data: (1) unlinked CIS data (each site evolves on its own own tree drawn randomly from the distribution of gene trees expected for the true simulation parameters under the MSC model), and (2) multilocus data (a sequence of length *l* is simulated for each locus on an underlying gene tree drawn randomly from the expected gene tree distribution). The simulations were performed as follows:

1. Use **ms** (Hudson, 2002) to generate *N* * *γ* gene tree samples under the MSC model based on the parental tree S1 and *N* * (1 − *γ*) gene tree samples based on the parental tree S2 in Figure 1;
2. Use **seq-gen** (Rambaut and Grass, 1997) to generate DNA sequences of length *l* for each gene tree under the specified nucleotide substitution model (*l* = 1 for CIS data);
3. Count the 15 site pattern frequencies;
4. Compute the test statistics for LRT, *HyDe* and ABBA-BABA methods, and test the hypothesis (1).
5. Repeat steps 1–4 *W* times to obtain type I error and statistical power for the three methods.

All steps in the simulations were carried out in the R statistical software (R Core Team, 2018). In step 1, time is measured in coalescent units (number of generations scaled by 2*N*_*e*_, where *N*_*e*_ is the effective population size), and we set the population size parameter *θ* = 4*N*_*e*_*μ* = 0.002 (constant throughout the tree), where *μ* is the mutation rate. For the speciation times in Figure 1, we assigned the vector (*τ*_1_, *τ*_2_, *τ*_3_) = *b*·(0.25, 0.5, 1.0), where different choices of *b* then involve stretching or shrinking the network; any choice of *b* results in trees that satisfy the molecular clock. We considered two choices for *b*: *b* = 1 and 2 to represent short and long branch trees, respectively. The hybridization parameter *γ* is chosen to be 0 or to vary from 0.06 to 0.5 by 0.02. For CIS data, we simulate *N* = 100*K*, 250*K*, 500*K* and 1*M* genes with one DNA site for each, while for multilocus data, we simulate *N* = 1*K*, 2.5*K*, 5*K* and 10*K* genes with length *l* = 100 in step 2. In that case, we have the same length of simulated DNA alignments for CIS and multilocus data. In steps 5, we chose *W* =500.

Though the LRT and *HyDe* are both derived under the JC69 model, DNA sequence may evolve under more complex models. To evaluate performance of the three methods under different substitution models, we studied two options for CIS data generation in step 2: JC69 and the general time reversible model (GTR) with mutation rates 1.0, 0.2, 2.5, 0.75, 3.2, 1.6 and base frequencies *π*_*A*_ = 0.22, *π*_*C*_ = 0.28, *π*_*G*_ = 0.22, *π*_*T*_ = 0.28.

### 2.4 Application to real data

#### 2.4.1 *Heliconius* Butterflies

DNA sequence data from Martin et al. (2013) were downloaded for four populations of *Heliconius* butterflies (248,822,400 sites; available on Dryad). We selected a single individual from four population each of which represents a distinct species: *Heliconius*

*melpomene rosina, H. m. timareta, H. cydno*, and the outgroup, *H. hecale*. Significant hybridization was detected in *H. cydno* at the population level in Martin et al. (2013) and using *HyDe* in Blischak et al. (2018) where they used multiple individuals per population. For counting the 15 site pattern frequencies, we only include sites with explicit nucleotides for all the species. Therefore, we have 128,321,514 sites for the four populations of *Heliconius* butterflies in the final analysis.

#### 2.4.2 *Sistrurus* Rattlesnakes

We applied our LRT to examine the question of whether populations of *Sistrurus catenatus* rattlesnakes in northwest and central Missouri are of hybrid origin. These populations are found to include individuals with morphological characteristics intermediate between *S. c. catenatus* and *S. c. tergeminus* (Gloyd, 1940; Evans and Gloyd, 1948). While a hypothesis of hybridization is plausible based on morphological similarity, it is also possible that this similarity is due to evolutionary or ecological factors. Gibbs et al. (2011) did not find evidence of hybridization between *S. c. catenatus* and *S. c. tergeminus* based on microsatellite and mitochondrial markers, indicating that the individuals in Missouri were *S. c. tergeminus*. Gerard et al. (2011) developed and used a likelihood ratio test based on observed gene tree distributions to analyze these data, and also found no genetic evidence of hybridization between *S. c. catenatus* and *S. c. tergeminus*.

We used the dataset of Gerard et al. (2011) to examine this question. The data consist of twelve genes (A, ATP, 1, 4, 11, 25, 31, 41, 61, 63, ETS, and GAPD) analyzed by Kubatko et al. (2011) (see their Table 2). The original dataset includes fourteen individuals: four of *S. c. catenatus*, four of *S. c. tergeminus*, four of the individuals in the putative hybrid zone, and two from outgroup populations of *Agkistrodon contortix* and *A. piscivorus*. We selected a single individulal from the three *Sistrurus* populations and one outgroup species. After excluding ambiguous sites, the dataset included 4 individuals and 7,663 sites.

## 3 Results

### 3.1 Simulation study

We plot test sizes for testing hybridization using the ABBA-BABA test, *HyDe*, and the LRT under JC69 for the short and long branch trees for different values of *γ* between 0 and 0.5. (see the Supplemental Material for figures under all of the simulation settings). As a representative example, Figures 2 and 3 show plots of the test results for the short and long branch trees with sequence length 100*K*. From these plots, we observe that all methods exhibit increased power as *γ* gets closer to 0.5, a pattern that becomes more prominent with an increase in dataset size. Comparing the three methods, we see that the LRT is more powerful in all cases. Even when the nucleotide substitution model is misspecified, type I error rates were reasonably controlled by the LRT for CIS data, while the ABBA-BABA test has inflated type I error in some cases (see Figure 3 and Supplemental Material Sections S1.1 and S1.2).

**Figure 2:**
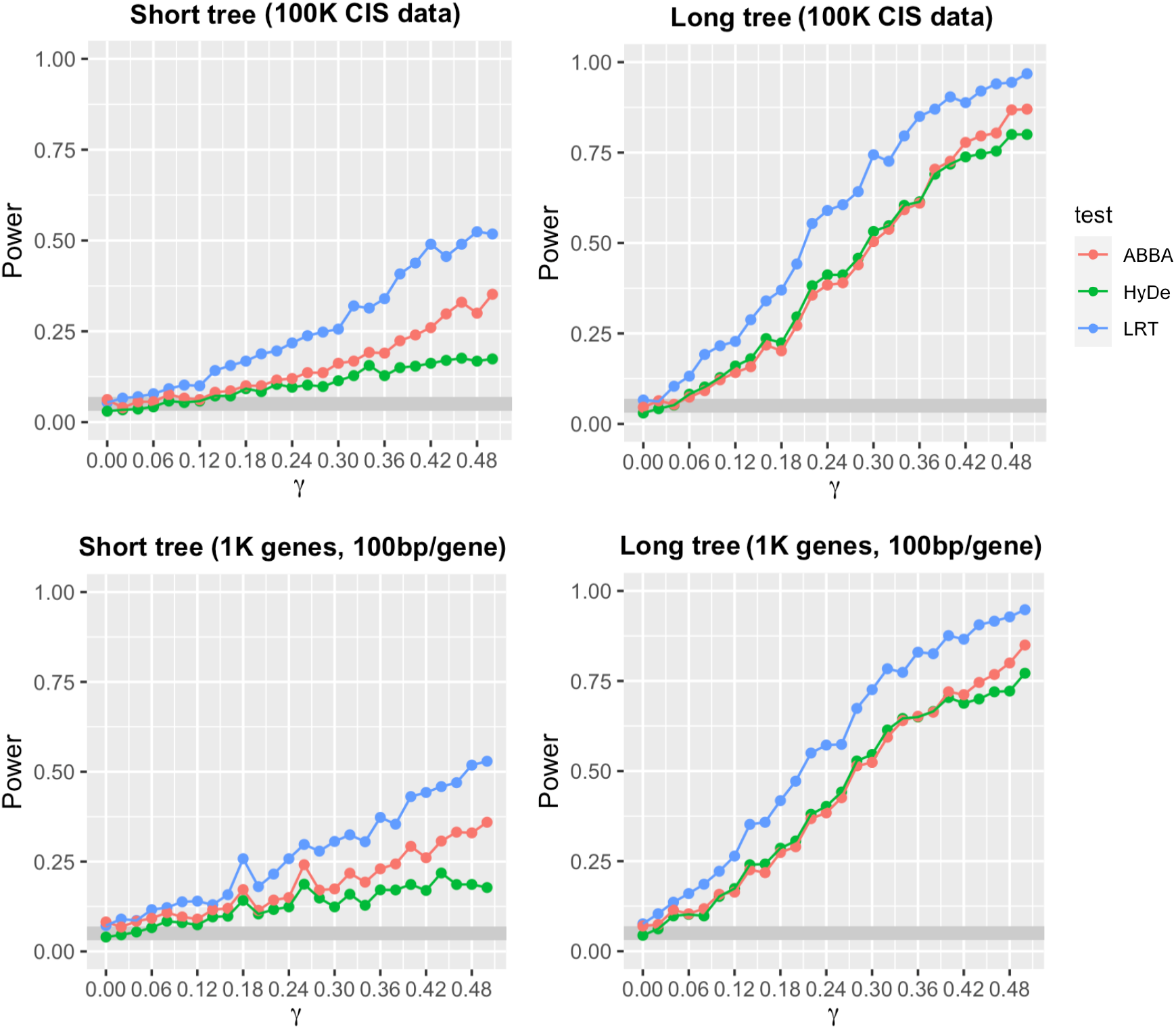
Hybrid detection power under JC69 in the ABBA-BABA test, *HyDe*, and the LRT for differing inheritance parameter values (*γ*) with sequence length 100K for the short and long branch tree. The top plots show test results from 500 Coalescent Independent Sites (CIS) datasets and the bottom plots show test results from 500 multilocus datasets. The shaded area gives the expected acceptance region of the empirical type I error rate in 500 simulation replicates.

**Figure 3:**
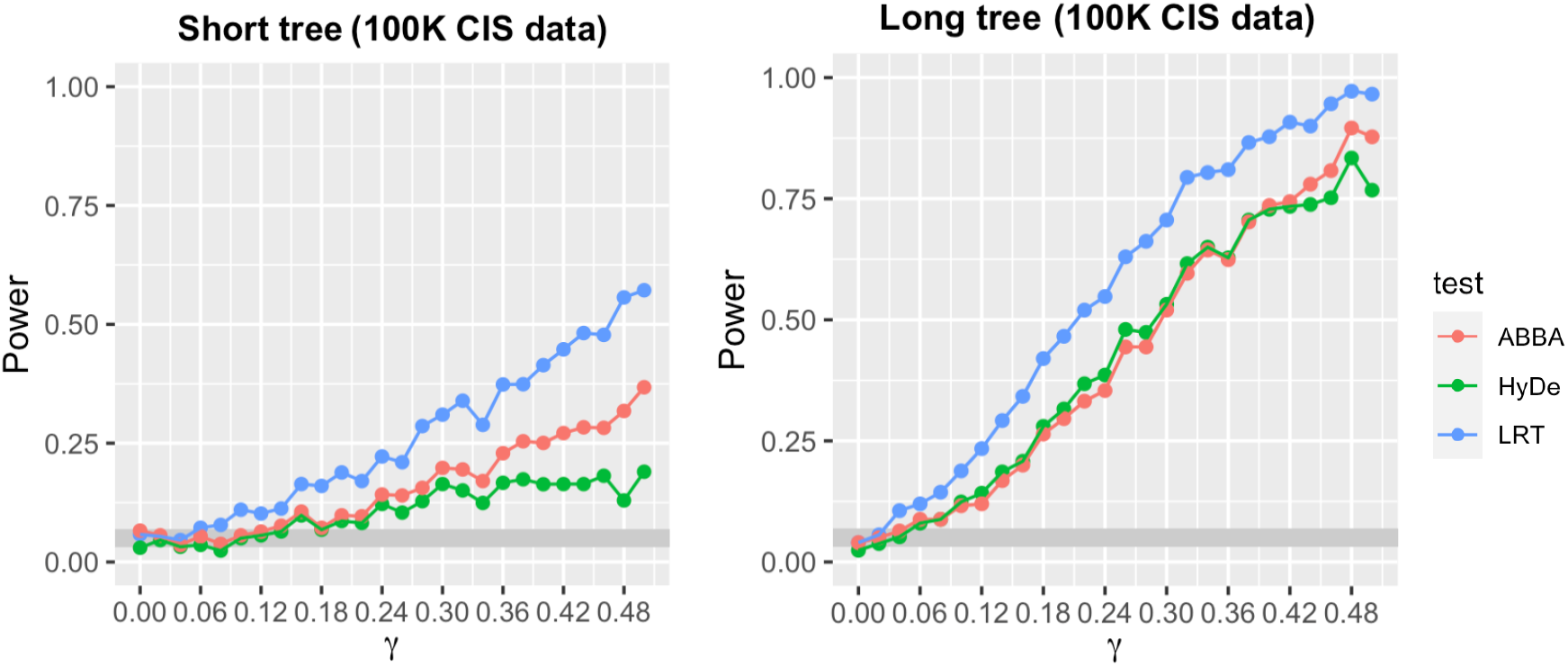
Hybrid detection power under GTR in the ABBA-BABA test, *HyDe*, and the LRT in different inheritance parameter values (*γ*) with sequence length 100K. The two plots present test results from 500 CIS datasets for the short and long branch tree, respectively. The shaded area gives the expected acceptance region of the empirical type I error rate in 500 simulation replicates.

For multilocus data, the type I error is a little inflated in most cases for the long branch tree and the two cases with smaller data sizes for the short branch tree (see Supplemental Material Sections S2.2 and S2.1, respectively.). In these cases, however, the ABBA-BABA test and *HyDe* have similar problems. This may be because the limited number of genes is not enough to fully characterize the gene tree distribution under the coalescent model. Although the statistical power is low overall for the short branch tree, we do not see much difference in the power or type I error for multilocus data with the same length for all methods in comparison with unlinked sites data. Similar patterns are observed when the number of loci is increased to 250K, 500K and 1M. As expected, the power for all three methods approaches 100% when the number of sites is 1M (see the Supplemental Material).

### 3.2 Empirical datasets

Table 1 shows the results of testing hybridization for the empirical data sets using the LRT, *HyDe* and the ABBA-BABA test. All three methods detect hybridization for the *Heliconius* data set with p-values less than 0.0001, consistent with previous work. For the *Sistrurus* rattlesnakes, all three methods fail to reject the null hypothesis of no hybridization at *α* = 0.05, again consistent with previous studies. Table 1 also shows the estimates of the inheritance parameter 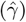 from *HyDe* and the LRT, which can be used to help decide if a network is preferable to a binary tree as a representation of the speciation history of a group. For example, the estimates of *γ* for the *Heliconius* data set are very close to 0.5 with small p-values, strongly suggesting a role for hybridization in the history of *H. cydno*. Conversely, for the *Sistrurus* data set, the estimates of *γ* are less than 0.1, suggesting little to no gene flow following speciation.

**Table 1:** Results of testing hybridization for the empirical datasets. 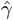 is the estimate of inheritance parameter *γ*.

| Datasets | # of sites | LRT |  | *HyDe* |  | ABBA-BABA |
| --- | --- | --- | --- | --- | --- | --- |
| | | p value | $\hat{\gamma}$ | p value | $\hat{\gamma}$ | p value |
| *Heliconius* butterflies | 128,321,514 | < 0.0001 | 0.493 | < 0.0001 | 0.415 | < 0.0001 |
| *Sistrurus* rattlesnakes | 7,663 | 0.064 | 0.066 | 0.380 | 0.036 | 0.382 |

## 4 Discussion

In this article, we develop the likelihood for a 4-taxon network under JC69 and the multispecies coalescent model. Based on that, we propose a likelihood ratio test for hybridization given a 4-tip species tree topology. Simulation studies demonstrate that our method achieves higher statistical power with reasonable type I error than the other two popular methods of testing hybridization, *HyDe* and the ABBA-BABA test, when sequence data are simulated under JC69. The increase in power is especially evident when the number of independent sites is limited and the species are recently diverged. Our simulations demonstrate that the test performs well for both coalescent independent sites and for multilocus data without much difference in power given the same overall sequence length. We used two empirical data sets to highlight the performance of the method in practice. We note that for all of the methods examined, the type I error may be inflated when the number of genes is limited. Thus, we encourage users to consider the estimate of *γ* when drawing conclusions. From the biological perspective, an estimated value of the hybridization parameter close to 0 is likely to indicate a lack of signal for hybridization. From a model selection perspective, we may not want to consider a network over a binary tree in such cases.

A primary innovation of our method is that we are more likely to detect the hybridization with less data than we are using current methods. In addition, the method provides maximum likelihood estimates (MLEs) for all the branch lengths, for the population size parameter, and for the hybridization parameter. That means the estimates share all the desirable statistical properties of MLEs, like consistency, asymptotically normality, and asymptotic efficiency. We are currently working to extend this method to larger networks.

The assumptions that (1) nucleotide sites evolve according to the JC69 substitution model and (2) effective population sizes are constant throughout the tree permit the use of formulas from Chifman and Kubatko (2015) for computing the 15 site pattern probabilities. Empirical data, however, may evolve under a nucleotide substitution model more complex than JC69. Our simulation studies indicate that our method is robust to data arising from the GTR model, though we did not exhaustively check all possible substitution rate matrices. When the true nucleotide substitution model differs greatly from JC69, our method may lead to inflated type I error. In these cases, the estimate of the hybridization parameter may provide a meaningful complement to the p-value in terms of biological interpretation. We are currently investigating approaches for extending our method to larger networks with multiple hybridization events. In that case, *HyDe* has multiple testing problems, while our method provides the possibility of using a likelihood-related score for model selection.

Code and functions that were used to carry out the simulation study and empirical analysis are presented in the Supplemental Material or can be obtained by contacting author Jing Peng at.

## 5 Additional

### 5.1 Funding

The authors have no funding to acknowledge.

### 5.2 Supporting Information

Supplemental Material is included.

