## Supplemental Material for "A Likelihood Ratio Test for Hybridization Under the Multispecies Coalescent"

June 6, 2023

#### S1. Comparison of three hybridization tests under GTR model

In this section, we show the supplemental figures of detection power and type I error using our LR test, *HyDe* and *ABBA-BABA* test under GTR model with different branch lengths and number of sites in section 2.3.

##### S1.1. Short branch tree

In this section, for the speciation times in Figure 1, we assigned the vector $(\tau_1, \tau_2, \tau_3) = (0.25, 0.5, 1.0)$. The hybridization parameter $\gamma$ is chosen to be 0 or to vary from 0.06 to 0.5 by 0.02.

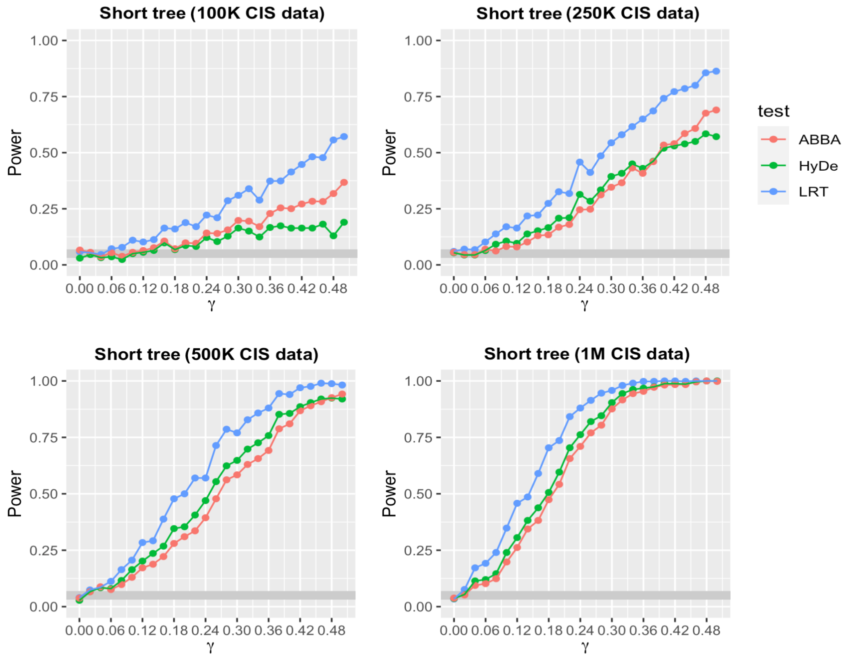


Figure 1: Hybrid detection power under the GTR model in the *ABBA-BABA* test, *HyDe*, and the LR test in different inheritance probabilities ($\gamma$) with 100K, 250K, 500K and 1M CIS sites, respectively. The shaded area is the expected acceptance region of the empirical type I error rate in 500 simulation replicates.

##### S1.2. Long branch tree

In this section, for the speciation times in Figure 1, we assigned the vector $(\tau_1, \tau_2, \tau_3) = (0.5, 1.0, 2.0)$. The hybridization parameter $\gamma$ is chosen to be 0 or to vary from 0.06 to 0.5 by 0.02.

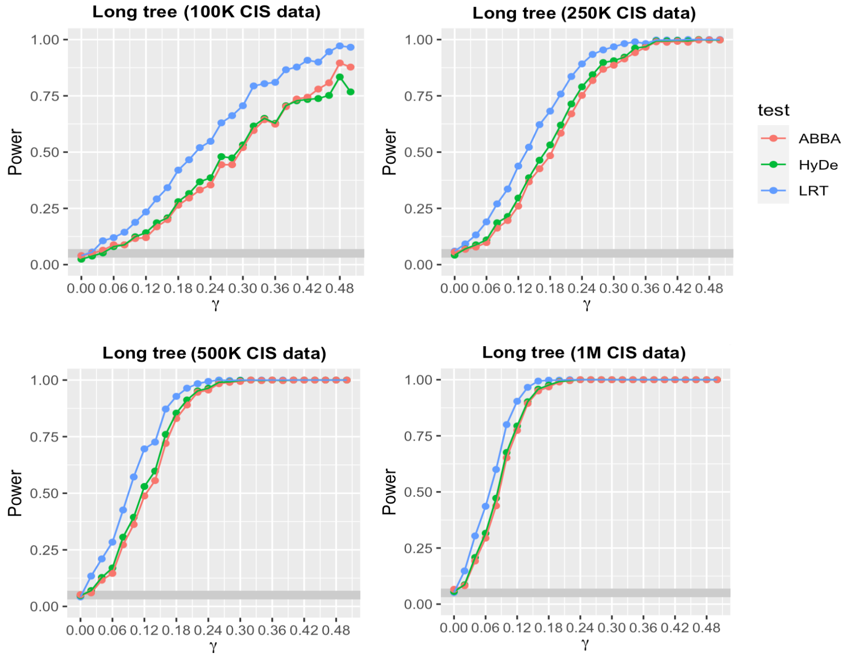


Figure 2: Hybrid detection power under the GTR model in the *ABBA-BABA* test, *HyDe*, and the LR test in different inheritance probabilities ($\gamma$) with 100K, 250K, 500K and 1M CIS sites, respectively. The shaded area is the expected acceptance region of the empirical type I error rate in 500 simulation replicates.

### S2. Comparison of three hybridization tests under JC69 model

In this section, we show the supplemental figures of detection power and type I error using our LR test, *HyDe* and *ABBA-BABA* test under GTR model with different branch lengths and number of sites in section 2.3.

#### S2.1. Short branch tree

In this section, we consider CIS and multilocus datasets. For the speciation times in Figure 1, we assigned the vector $(\tau_1, \tau_2, \tau_3) = (0.25, 0.5, 1.0)$. The hybridization parameter $\gamma$ is chosen to be 0 or to vary from 0.06 to 0.5 by 0.02.

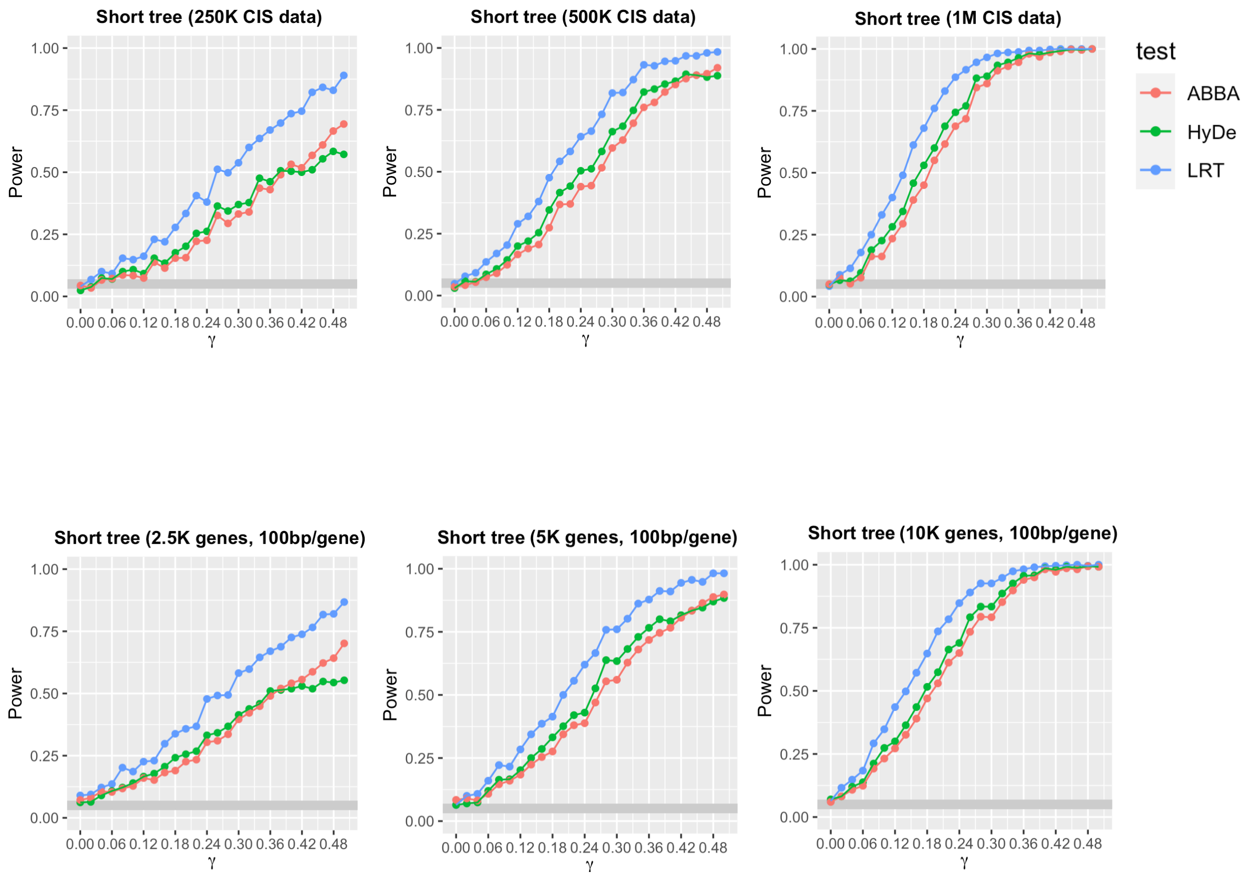


Figure 3: Hybrid detection power under JC69 in the *ABBA-BABA* test, *HyDe*, and the LR test in different parental contributions ($\gamma$) with seuqence length 250K, 500K and 1M CIS sites, respectively. The shaded area gives the expected acceptance region of the empirical type I error rate in 500 simulation replicates.

#### S2.2. Long branch tree

In this section, we consider CIS and multilocus datasets. For the speciation times in Figure 1, we assigned the vector $(\tau_1, \tau_2, \tau_3) = (0.5, 1.0, 2.0)$. The hybridization parameter $\gamma$ is chosen to be 0 or to vary from 0.06 to 0.5 by 0.02.

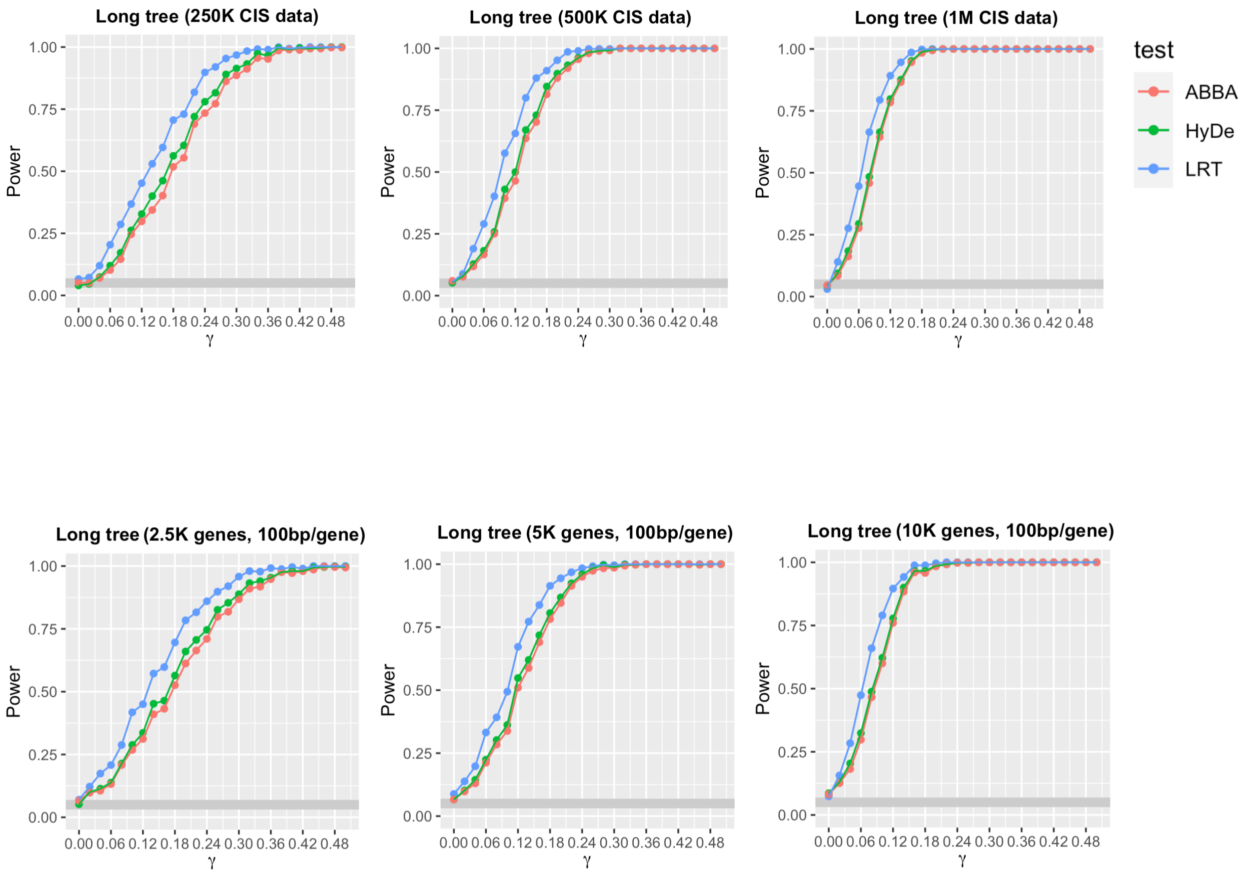


Figure 4: Hybrid detection power under JC69 in the *ABBA-BABA* test, *HyDe*, and the LR test in different parental contributions ($\gamma$) with seuqence length 250K, 500K and 1M CIS sites, respectively. The shaded area gives the expected acceptance region of the empirical type I error rate in 500 simulation replicates.
